# Improving the value utilization of tuna peptide powder for the cosmetics field through ozone oxidation

**DOI:** 10.1101/2024.08.18.608506

**Authors:** Haolong Zheng, Shiyang Gu, Shiqi Huang, Yan Zhang, Feng Xu, Daofei Lv, Wenbing Yuan, Kongyu Zhu, Xin Chen

## Abstract

To improve the utilization efficiency of small molecule peptides in marine proteins and promote their application in cosmetics, this study adopts ozone oxidation to conduct deodorization research on tuna small molecule peptides. To devise a method for the ozone oxidation and deodorization of tuna peptide powder suitable for the cosmetics field, orthogonal tests were carried out. The time, ozone concentration, and temperature were adjusted to determine the optimal conditions for ozonation and deodorization of tuna peptide powder. The results showed that at 50°C and an ozone concentration of 99.10 × 103 mg/m^3^, wet ozone oxidation for 40 min can also significantly reduce the fishy smell. Under these conditions, the color of tuna peptide powder treated with wet ozone was lighter. In addition, the main volatile compounds were detected by gas chromatography–mass spectrometry (GC-MS). The content of the main fishy volatile substance, hexanal, decreased by 66.4%, and two octyl aldehydes were observed. The results also showed that the protein activity of tuna peptide powder after ozonation remained unchanged. Thus, this study presents an approach to the ozone oxidation of tuna peptide powder that is suitable for cosmetics.

## INTRODUCTION

Tuna (Perciformes, Scombridae) is a type of fish that lives mainly in open oceanic waters at mid and low latitudes. It is popular among a wide range of consumer groups due to its high economic and nutritional value^[1][2][3]^. Tuna is rich in high-quality protein and n-3 fatty acids, such as docosahexaenoic acid, eicosapentaenoic acid, and other essential nutrients, which play a vital role in disease prevention and treatment^[3]^. In recent years, foods, cosmetics, and healthcare products made from tuna have become popular among consumers because of the high protein content, nutritional value, and low fat content of the fish.

Marine biological peptides are natural active peptides extracted from marine organisms, possessing a diverse range of physiological functions and benefits. According to Amnuaikit et al.^[4]^, cosmetic products containing marine biological peptides have the ability to enhance facial skin moisture, minimize pores and skin wrinkles, and significantly brighten the skin when used regularly for a minimum of two weeks. It is 100% absorbable through the skin.

However, the fishy odour of tuna peptide powder does not allow it to be used directly in cosmetics. Fishy odor is one of the major factors affecting the quality of fish and their acceptance by consumers^[5][6][7]^. As a result, effective deodorization is a major challenge for the industrial reprocessing of fish products. Promoting the high-value utilization of marine products through effective deodorization is crucial, as it facilitates their utilization in cosmetics, health products, and food products. At present, the main methods of deodorization used are as follows:

Masking is one of the earliest methods, which uses the volatile gas of other substances to cover the smell of the fish itself^[1]^. However, masking does not completely remove the original odor and introduces new substances. Zeng et al. adsorbed tilapia enzymolysis solution with a selective adsorbent, to achieve effective fish odor removal and a relatively high protein recovery rate^[8]^. However, this absorbent was responsive to different removal gases. The choice of adsorbent is limited, and the high-cost saturated adsorbent also needs follow-up treatment to prevent environmental pollution and adverse effects. When Wu et al. processed eel meat, the addition of 1.5% cyclodextrin β significantly removed the odor of eel meat after stirring for 90 min^[9]^. However, this method cannot completely remove the volatile odor, and the embedding effect of macromolecular volatile substances is poor. Taking silver carp as the treatment object, Fu et al. adopted an alkali treatment^[10]^. After treatment, the content of ichthyoid in the water of silver carp was greatly reduced, but this method destroys the nutrients of the aquatic product itself. The basicity of the sample cannot easily be removed, and the final waste liquid needs to be treated before discharge. In biological deodorization, different types of microorganisms are added at a certain temperature and humidity, and microbial metabolites are used to reduce the concentration of odorous substances. Xu et al. fermented silver carp surimi with yeast at 35°C for 1.5 h^[11]^. The fishy taste of the silver carp was effectively reduced, but this method has poor applicability. Different fish substances have different resistances to different microorganisms, and other substances are generated via microbial activity due to the long treatment period.

We found that the most suitable method for deodorization of polymer active peptide components in cosmetics is ozone oxidation. Ozone is a strong oxidizing agent that may be more inclined to produce highly oxidizing hydroxyl radicals in water that can oxidize reducing substances and decompose organic matter in aquatic products. Ozone reacts with aldehydes, alcohols, amines, and other volatile substances in aquatic products to reduce or eliminate fishy odor substances and achieve deodorization^[2][12]^. The volatile component content of silver carp minced meat treated with floating ozone technology decreased without introducing other odors^[13]^. In contrast, the use of ozone oxidation technology for removing the fishy odor of tuna peptide powder offers several advantages. (1) Efficiency: Compared to other techniques, ozone oxidation exhibits a higher treatment efficiency and better results than traditional methods. (2) No harmful effects on humans: Ozone oxidation treatment produces no pollutants or residues, ultimately generating only water and oxygen. This ensures there is no secondary pollution harmful to human body or the environment. The reactions are as follows: O_3_ + H_2_O → H_2_O_2_ + O_2,_ 5H_2_O_2_ + O_3_ → 4O_2_ + 5H_2_O^[14]^. (3) Accessibility of equipment: Ozone oxidation equipment and technology are relatively simple and can be easily implemented, maintained, and automated.

Therefore, the purpose of this study is to utilize ozone oxidation to create a deodorizing process suitable for tuna polymer peptide powder in the field of cosmetics. After deodorization, the tuna peptide was assessed by sensory evaluation, gas chromatography–mass spectrometry (GC-MS), cold/heat stability analysis, determination of amino nitrogen and total nitrogen content before and after oxidation, and other methods to confirm the comprehensive feasibility of the process conditions. The results of this study will provide a theoretical basis and insight for the development of ozone deodorization technology for seafood and have practical significance for the application of this technology in cosmetics.

## MATERIALS AND METHODS

The materials used in this study include Pulvis tuna oligopeptide powder (Liaoning Taiai Peptide Bioengineering Technology Co., Ltd., Dalian, China) and sodium chloride (AR; Aladdin Reagent (Shanghai) Co., Ltd., Shanghai, China). Instruments used include a Kjeldahl nitrogen analyzer (K9840, Hanon Advanced Technology Group Co., Ltd., Jining, China), a freeze dryer (LGJ-10, Sihuan Keyi Technology Development Hebei Co., Ltd., Tangshan, China), an ozone generator (QD-Y, Guangzhou Qida Environmental Protection Equipment Co., Ltd., Guangzhou, China), a heat-collecting thermostatic magnetic stirrer (DF-101S, Gongyi Yuhua Instrument Co., Ltd.), and ozone concentration detectors (Model 106-H and 202-L, Passkey Technology Co., Ltd., Boulder, USA).

### Ozone Concentration Detection

As van Leeuwen emphasized^[15]^, results obtained without specifying the ozone dose are inherently meaningless. To ensure accuracy in our experiment, ozone concentration detectors were used to obtain the ozone concentration^[16]^.

### Dry Ozonation of Tuna Peptides

First, 1 g of tuna peptide powder ( ± 0.05 g) was weighed using an electronic balance. The measured powder was then placed in a conical flask equipped with two tubes and fitted with a rubber stopper. One tube was connected to the ozone generator and the other to a water tank designated for exhaust gas treatment (Figure 1). After connecting the ozone generator water supply device, the main power was turned on the device was allowed to run for 5 min. Subsequently, the ozone switch was turned on and set to deliver specific ozone concentrations (0.1 mg/L–104.6 mg/L) over a specified period (0–300 min). A consistent ozone flow rate of 1.5 L/min was maintained throughout all experiments. After the ozone supply was switched off, the rubber stopper was removed, and the mixture was allowed to stand for 10 min. When the ozone in the flask had decomposed, the peptide powder was removed for sensory evaluation.

**FIGURE 1.**
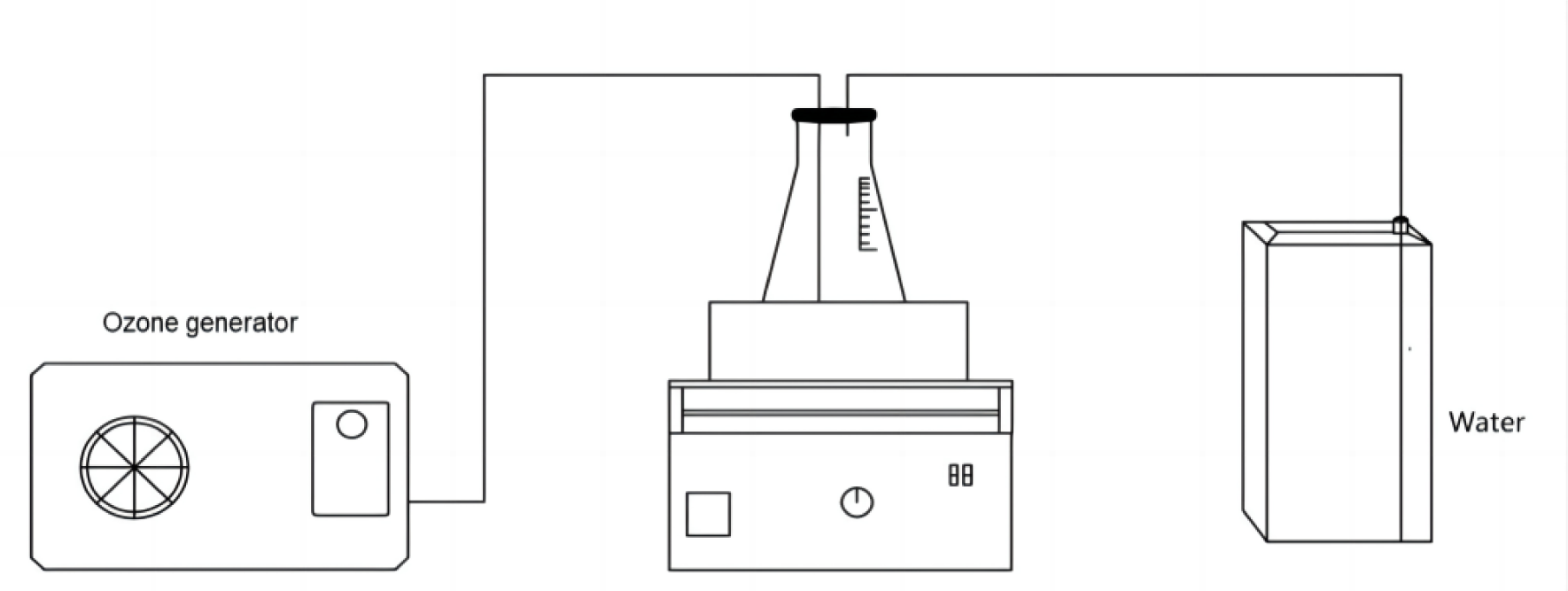
Apparatus for the dry ozonation of tuna peptides.

### Wet Ozonation of Tuna Peptides

First, an electronic balance was used to weigh 1 g of tuna peptide powder. Next, a graduated cylinder was used to measure 5 mL of distilled water, which was placed in a three-necked flask. Then, the weighed tuna peptide powder was placed in the three-necked flask containing distilled water, and it was ensured that the rubber stoppers were securely in place on each neck. Then, a tube was inserted through the stoppers on both sides, connecting one end to the ozone generator and the other to the tap water tank for exhaust gas treatment (Figure 2). After connecting the ozone generator water supply device, the main power of the ozone generator was switched on, and it was allowed to run for 5 minutes before the ozone switch was activated and set to provide specific ozone concentrations (0.1 mg/L–104.6 mg/L) over a specified time (0–300 min). A consistent ozone flow rate of 1.5 L/min was maintained throughout all experiments. After the ozone supply was switched off, the rubber stopper was removed, and the mixture was allowed to stand for 10 min. When the ozone in the flask had decomposed, the peptide powder solution was removed and freeze-dried to obtain a dry peptide powder for sensory evaluation.

**FIGURE 2.**
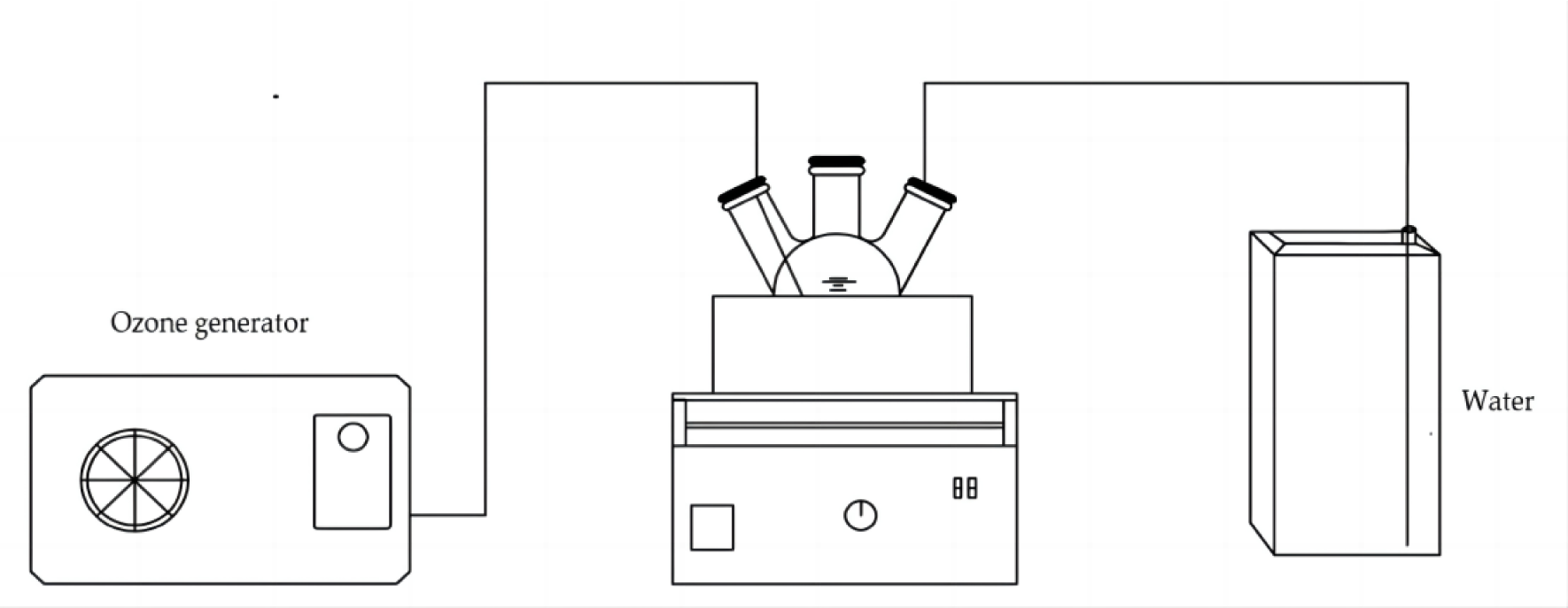
Apparatus for the wet ozonation of tuna peptides.

### Sensory Evaluation

With reference to Khazandi et al.^[17]^, the filtrate was sensorily evaluated using a 5-point scale (1, no fishy odor; 2, slight fishy odor; 3, moderate fishy odor; 4, heavy fishy odor; 5, very heavy fishy odor). Deodorized tuna was sensorily evaluated using a “fishy score,” which was calculated by summing the weighted scores for fishy odor and fishy taste. The higher the scores, the stronger the fishy odor and fishy taste^[18][19]^. Fishy odor and taste were scored using a five-point scale: 1, no fishy odor/taste; 2, slight fishy odor/taste; 3, moderate fishy odor/taste; 4, strong fishy odor/taste; 5, very strong fishy odor/taste. The overall average fishy score V for each sample was calculated as follows:

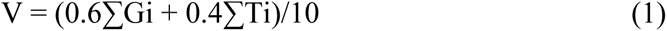

where Gi is the fishy odor score assigned to the sample by the ith sensory evaluator, Ti is the fishy taste score assigned to the sample by the ith sensory evaluator, and 10 was the number of evaluators.

Sensory evaluation was repeated on days 1, 3, and 5 following the above method to obtain three sensory scores (Figs. 3–5).

**FIGURE 3.**
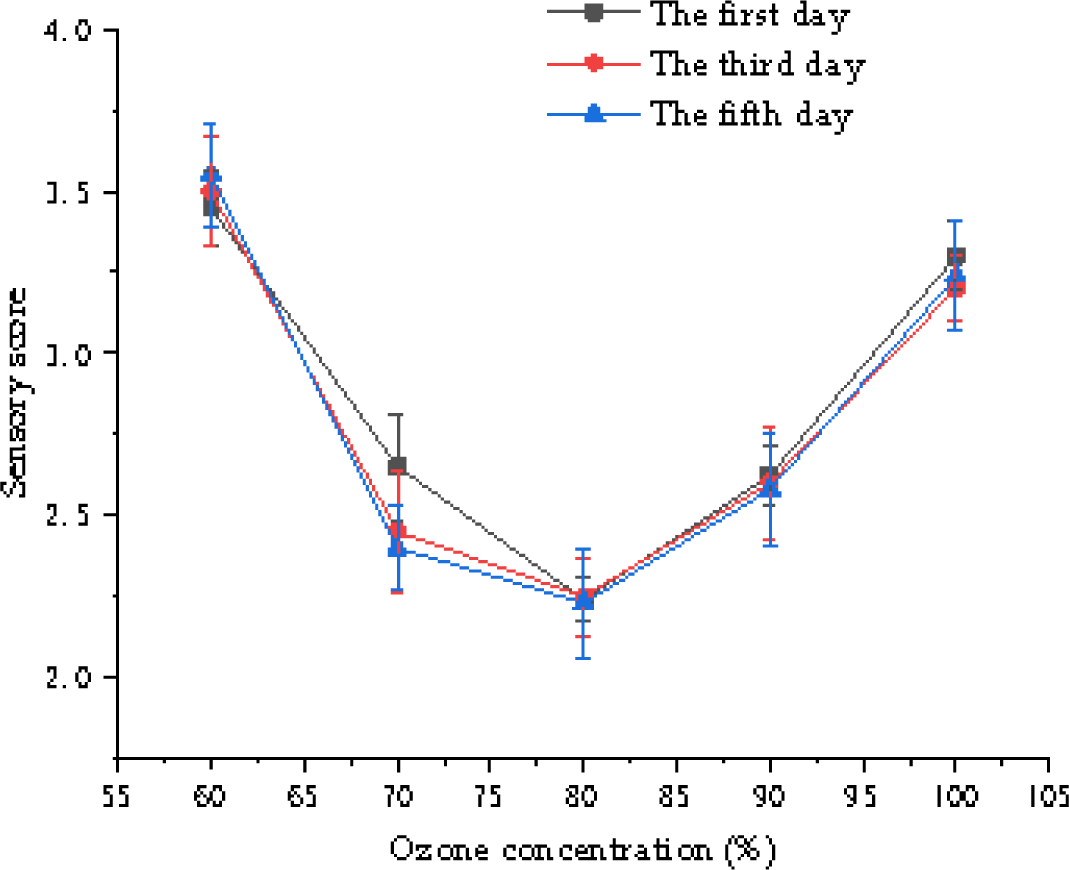
Effect of ozone concentration on sensory score.

**FIGURE 4.**
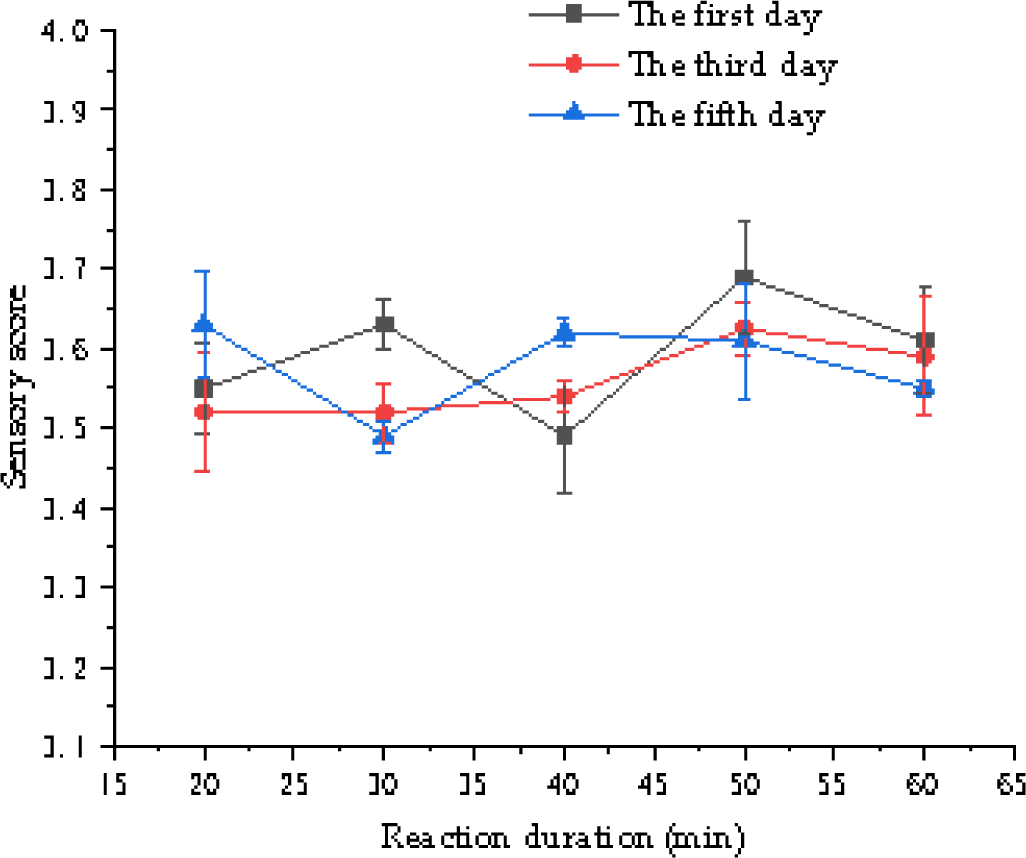
Effect of reaction time on sensory score.

**FIGURE 5.**
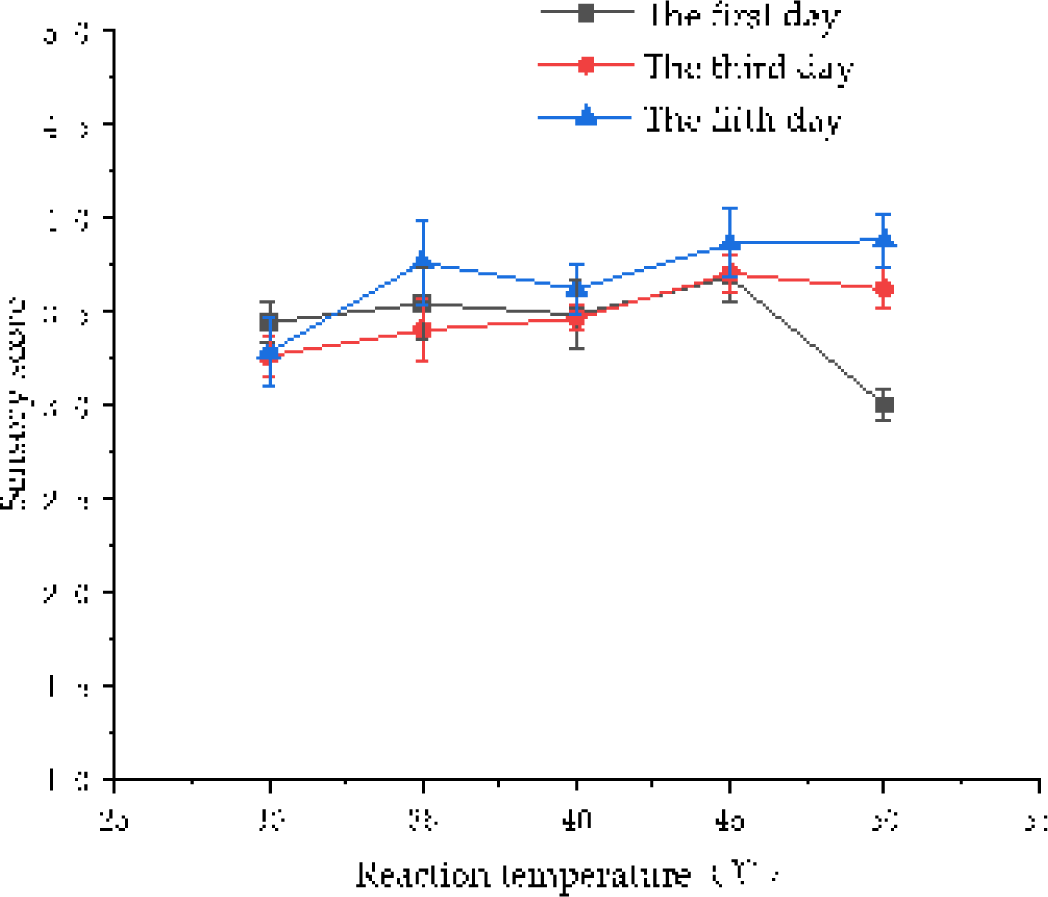
Effect of reaction temperature on sensory score.

### Determination of Total Nitrogen and Amino Nitrogen

The total nitrogen and amino nitrogen contained in the samples were respectively determined according to the GB5009.5-2016 and GB5009.235-2016 national standards of the People’s Republic of China.

### Long-term Low-/High-Temperature Stability Tests

Temperature stability tests were carried out on hand cream samples containing 1% ozone-treated tuna peptides. Each sample was separated into two vials (A and B), and each vial was sealed with a clean stopper. Vial A was subjected to a 3-day heating–cooling cycle that was repeated 10 times: the sample was placed in an incubator for 24 h at 40°C, left at room temperature for another 24 h, and finally placed in a refrigerator at –10°C for another 24 h. Vial A was observed for oil–water stratification and color changes during this process by visual comparison with vial B^[20]^.

### Test Conditions for GC-MS Analysis

The tuna peptide powder with the best sensory evaluation after ozonation was selected for determination by GC-MS.

Pre-treatment method: Headspace solid-phase microextraction. First, 0.2 g of samples, 1.44 g of solid NaCl, and 8 mL of deionized water were sequentially added and mixed in a beaker. The mixture was then transferred to a headspace sampling bottle for sealing after dissolution. The bottle was placed in a magnetic agitator at 60°C and heated to equilibrium for 15 min, followed by adsorption with an activated SPEM extractor head for 40 min. The extractor head was placed at the injection port for analysis for 5 min.

GC conditions: An HP-5MS capillary column (30 m × 0.25 mm × 0.25 μm) was used, and the temperature of the KC interface was maintained at 240°C. He (99.99%) was used as the carrier gas at a flow rate of 1.0 mL/min without diverting. The initial temperature of the column was set at 40°C (Zhang et al. 2012), and the holding time was 2 min. The temperature of the machine was raised to 160°C at 4°C/min, increased to 250°C at 8°C/min, and maintained for 10 min.

MS conditions: Solvent removal time, 1 min; ion source, electron bombardment source (EI); ion source temperature, 230°C; ionization voltage, 70 eV; scanning mass range, 30–500 m/z; interface temperature, 280°C; four-stage bar temperature, 150°C.

## Results

### Dry Ozonation of Tuna Peptide Powder

Tuna peptide powder was subjected to 12 single-factor experiments involving dry ozonation, and the results of the sensory evaluation (fishy odor/taste) are shown in Table 1. The duration of dry ozonation had a small impact on the deodorization of the samples, while the ozone concentration significantly affected the sensory scores. The average sensory score was 4.5 at ozone concentrations of 0.1 mg/L–50.2 mg/L, which decreased to 3.7 when the ozone concentration was increased to 99.1 mg/L and decreased further to 3.5 at a concentration of 104.6 mg/L. These results are consistent with the finding by Wang et al. ^[19]^ that ozonated water had the best deodorizing effect on minced silver carp at an initial concentration of 0.78 mg/L. Therefore, increasing the ozone concentration effectively improved the deodorizing effect of dry ozonation.

**Table 1.** Sensory scores for samples in single-factor dry ozonation experiments.

| Experiment | Conditions* | Sensory score |
| --- | --- | --- |
| 1 | 30 min/50.2 mg/L | 4.7 ± 0.16 |
| 2 | 60 min/50.2 mg/L | 4.7 ± 0.15 |
| 3 | 90 min/50.2 mg/L | 4.5 ± 0.12 |
| 4 | 120 min/50.2 mg/L | 4.1 ± 0.16 |
| 5 | 180 min/50.2 mg/L | 4.1 ± 0.13 |
| 6 | 240 min/50.2 mg/L | 4.0 ± 0.14 |
| 7 | 300 min/50.2 mg/L | 4.0 ± 0.14 |
| 8 | 60 min/0.1 mg/L | 5.0 ± 0.24 |
| 9 | 60 min/4.2 mg/L | 4.8 ± 0.32 |
| 10 | 60 min/50.2 mg/L | 4.5 ± 0.26 |
| 11 | 60 min/99.1 mg/L | 3.7 ± 0.32 |
| 12 | 60 min/104.6 mg/L | 3.5 ± 0.40 |

\*Duration of ozonation/ozone concentration. The temperature is not given as it had a negligible effect.

The results obtained in the single-factor experiments were used to design nine orthogonal experiments on dry ozonation, and the sensory evaluation results (fishy odor/taste) are shown in Table 2. The optimum conditions for the dry ozonation of tuna peptide powder were found to be an ozone concentration of 99.1 mg/L and a treatment time of 180 min.

**Table 2.**
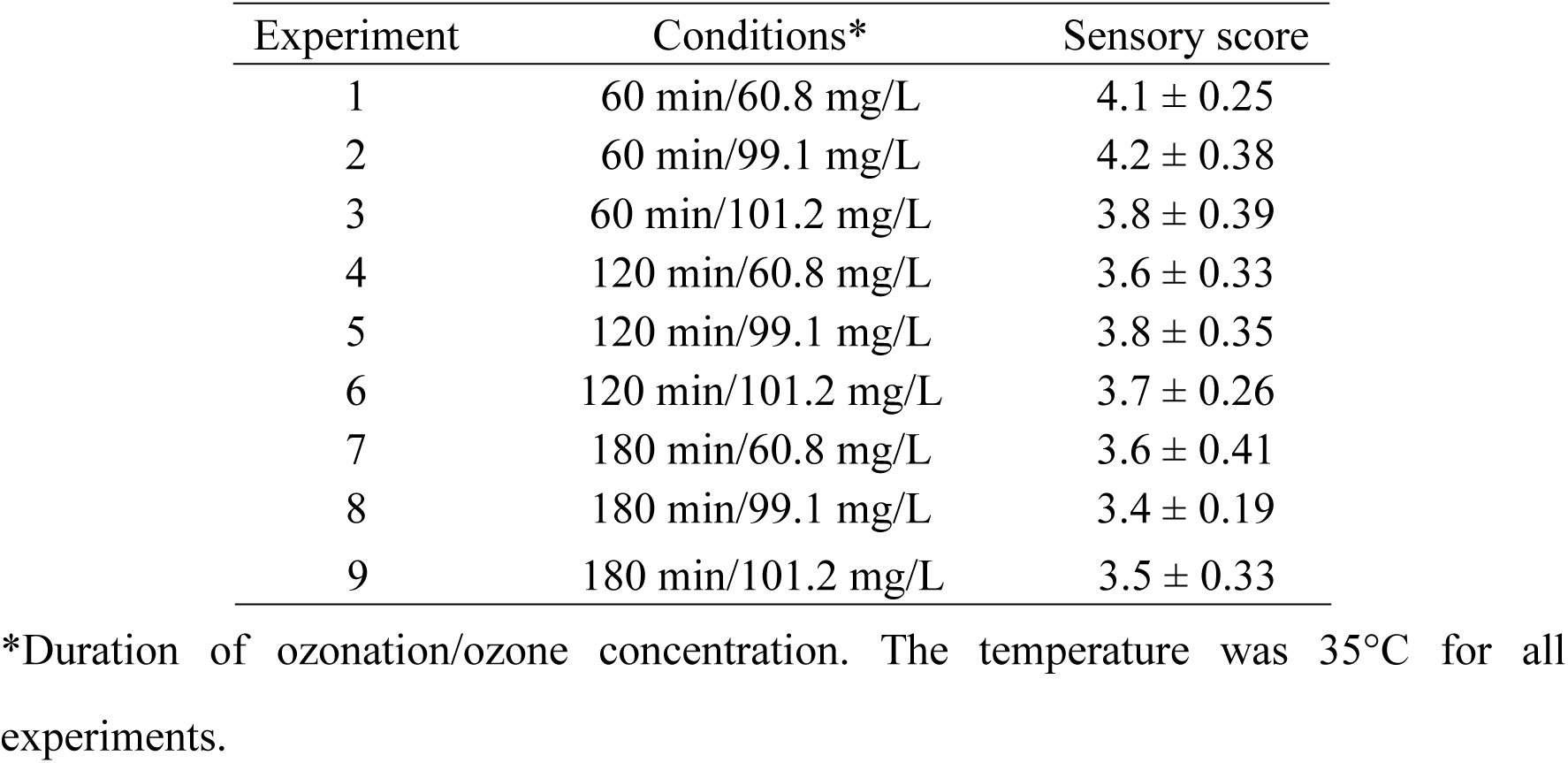
Sensory scores for samples in the orthogonal dry ozonation experiments.

### Wet Ozonation of Tuna Peptide Powder

Tuna peptide powder was treated by wet ozonation in 17 single-factor experiments, and the sensory evaluation results (fishy odor/taste) are shown in Table 3. When the reaction time was gradually increased from 30 to 300 min with the same ozone concentration and reaction temperature (experiments 1–7), the sensory scores ranged between 4.3 and 4.6. When the ozone concentration was gradually increased from 0.1 mg/L to 104.6 mg/L with the same reaction time and temperature (experiments 8–12), the sensory scores ranged from 2.2 to 5.0. When the reaction temperature was gradually increased from 30 to 70°C with the same reaction time and ozone concentration (experiments 13–17), the lowest sensory score was 4.0 and the highest was 4.4. Therefore, the reaction time and temperature had only a small impact on the deodorizing effect of wet ozonation, while the ozone concentration had a much larger influence.

**Table 3.** Sensory scores for samples in the single-factor wet ozonation experiments.

| Experiment | Conditions* | Sensory score |
| --- | --- | --- |
| 1 | 30 min/50.2 mg/L/35°C | 4.6 ± 0.13 |
| 2 | 60 min/50.2 mg/L/35°C | 4.6 ± 0.11 |
| 3 | 90 min/50.2 mg/L/35°C | 4.5 ± 0.14 |
| 4 | 120 min/50.2 mg/L/35°C | 4.3 ± 0.17 |
| 5 | 180 min/50.2 mg/L/35°C | 4.3 ± 0.08 |
| 6 | 240 min/50.2 mg/L/35°C | 4.4 ± 0.14 |
| 7 | 300 min/50.2 mg/L/35°C | 4.5 ± 0.11 |
| 8 | 60 min/0.1 mg/L/35°C | 5.0 ± 0.33 |
| 9 | 60 min/4.2 mg/L/35°C | 4.8 ± 0.32 |
| 10 | 60 min/50.2 mg/L/35°C | 4.5 ± 0.26 |
| 11 | 60 min/99.1 mg/L/35°C | 2.2 ± 0.27 |
| 12 | 60 min/104.6 mg/L/35°C | 2.8 ± 0.24 |
| 13 | 60 min/50.2 mg/L/30°C | 4.2 ± 0.23 |
| 14 | 60 min/50.2 mg/L/40°C | 4.2 ± 0.09 |
| 15 | 60 min/50.2 mg/L/50°C | 4.0 ± 0.22 |
| 16 | 60 min/50.2 mg/L/60°C | 4.2 ± 0.21 |
| 17 | 60 min/50.2 mg/L/70°C | 4.4 ± 0.26 |

\*Duration of ozonation/ozone concentration/temperature.

The results obtained in the single-factor experiments were used to design 25 orthogonal experiments of wet ozonation. The single-factor experiments revealed that reaction time and temperature had only a small impact on the deodorizing effect of wet ozonation. Therefore, these two variables were varied by only small amounts in the orthogonal experiments^[19]^. Specifically, reaction time was changed in increments of 10 min (i.e., 20, 30, 40, 50, and 60 min), while the reaction temperature was adjusted in increments of 5°C (i.e., 30, 35, 40, 45, and 50°C). As the ozone concentration had the greatest impact on the deodorizing effect of wet ozonation, this factor was altered in large increments (i.e., 50.2 mg/L, 60.8 mg/L, 99.1 mg/L, 101.2 mg/L, and 104.6 mg/L). The results obtained from the wet ozonation orthogonal experiments (Table IV) were then further analyzed^[21]^. Figures 3–5 respectively show plots of the sensory scores from the 25 orthogonal experiments against ozone concentration, reaction time, and reaction temperature.

The sensory score first decreased and then increased with increasing ozone concentration (Figure 3). When the ozone gas concentration was increased (the intake flow rate remaining unchanged), the concentration of ozone dissolved in the peptide powder solution after a short period increased, thereby increasing the mass transfer efficiency. The sensory score was the lowest (2.2) at an ozone concentration of 99.1 mg/L and then increased again upon further increasing the ozone concentration. The peptide powder solution had reached saturation at an ozone concentration of 99.1 mg/L. When the ozone concentration increased beyond 99.1 mg/L, excess ozone would introduce other odors, reducing the deodorizing effect.

The sensory score changed only slightly with increasing reaction time, producing a curve that approximates a straight line (Figure 4). Ozonation is a rapid process that can be completed in a short time due to the high oxidation–reduction potential (2.07 V) of ozone. Therefore, the reaction time had only a small impact on the deodorizing effect of wet ozonation^[22]^.

The sensory score changed only slightly with increasing reaction temperature, also producing a curve that approximates a straight line (Figure 5). Although the increase in the temperature of the peptide powder solution increased the activity of organic molecules, it also decreased the solubility of ozone and thereby reduced the mass transfer efficiency. As a result, the impact of increasing temperature on the deodorizing effect was reduced. Therefore, the reaction temperature also had only a small effect on the deodorizing effect of wet ozonation^[23]^.

The data presented in Table 4 and Figures 3–5 show that, of the three variables, ozone concentration has the greatest impact on the deodorizing effect of wet ozonation of tuna peptide powder, with reaction temperature and reaction time having only a small impact. The optimal conditions for treating tuna peptides by wet ozonation were as follows: ozone concentration, 99.1 mg/L; reaction time, 40 min; reaction temperature, 50°C. Under these conditions, the lowest sensory score of 2.3 was obtained, representing a decrease of 54%, which is similar to the largest decrease of 50.2 mg/L achieved by Chen et al.^[24]^ when deodorizing sea bass meat using a combined method and that of 46% achieved by Wang et al.^[17]^ when deodorizing minced silver carp using ozone water. The results obtained thus suggest that wet ozonation is effective in deodorizing tuna peptides.

**Table 4.** Sensory scores for the samples in the orthogonal wet ozonation experiments.

| Experiment | Conditions* | Sensory score |
| --- | --- | --- |
| 1 | 20 min/50.2 mg/L/30°C | 4.6 ± 0.26 |
| 2 | 30 min/50.2 mg/L/35°C | 4.4 ± 0.31 |
| 3 | 40 min/50.2 mg/L/40°C | 4.7 ± 0.25 |
| 4 | 50 min/50.2 mg/L/45°C | 4.3 ± 0.34 |
| 5 | 60 min/50.2 mg/L/50°C | 4.4 ± 0.33 |
| 6 | 20 min/60.8 mg/L/35°C | 3.7 ± 0.27 |
| 7 | 30 min/60.8 mg/L/40°C | 3.6 ± 0.11 |
| 8 | 40 min/60.8 mg/L/45°C | 3.6 ± 0.07 |
| 9 | 50 min/60.8 mg/L/50°C | 3.5 ± 0.43 |
| 10 | 60 min/60.8 mg/L/30°C | 3.6 ± 0.50 |
| 11 | 20 min/99.1 mg/L/40°C | 3.4 ± 0.23 |
| 12 | 30 min/99.1 mg/L/45°C | 2.9 ± 0.22 |
| 13 | 40 min/99.1 mg/L/50°C | 2.3 ± 0.31 |
| 14 | 50 min/99.1 mg/L/30°C | 2.6 ± 0.26 |
| 15 | 60 min/99.1 mg/L/35°C | 2.8 ± 0.27 |
| 16 | 20 min/101.2 mg/L/45°C | 3.1 ± 0.29 |
| 17 | 30 min/101.2 mg/L/50°C | 2.8 ± 0.31 |
| 18 | 40 min/101.2 mg/L/30°C | 2.9 ± 0.32 |
| 19 | 50 min/101.2 mg/L/35°C | 2.7 ± 0.33 |
| 20 | 60 min/101.2 mg/L/40°C | 3.1 ± 0.34 |
| 21 | 20 min/104.6 mg/L/50°C | 3.2 ± 0.23 |
| 22 | 30 min/104.6 mg/L/30°C | 2.8 ± 0.43 |
| 23 | 40 min/104.6 mg/L/35°C | 2.8 ± 0.53 |
| 24 | 50 min/104.6 mg/L/40°C | 2.7 ± 0.42 |
| 25 | 60 min/104.6 mg/L/45°C | 2.7 ± 0.43 |

\*Duration of ozonation/ozone concentration/temperature.

### Determination of Total Nitrogen and Amino Nitrogen in Ozone-treated Tuna Peptide Powder

As shown in Table 5, ozonation had a small impact on the total nitrogen and amino nitrogen content of tuna peptide powder. Compared with the original sample, the largest reductions in total nitrogen and amino nitrogen in the three test samples were 7.1% and 12.0%, respectively. These changes in total nitrogen and amino nitrogen were small compared with the previously reported nitrogen loss of 22.9% in treated Antarctic krill meal^[25]^. Therefore, nutrient-rich tuna peptide powder has the potential to be a useful source of collagen for cosmetic use, especially in applications where the use of collagen from cattle and pigs, the traditionally predominant sources of collagen, is limited.

**Table 5.**
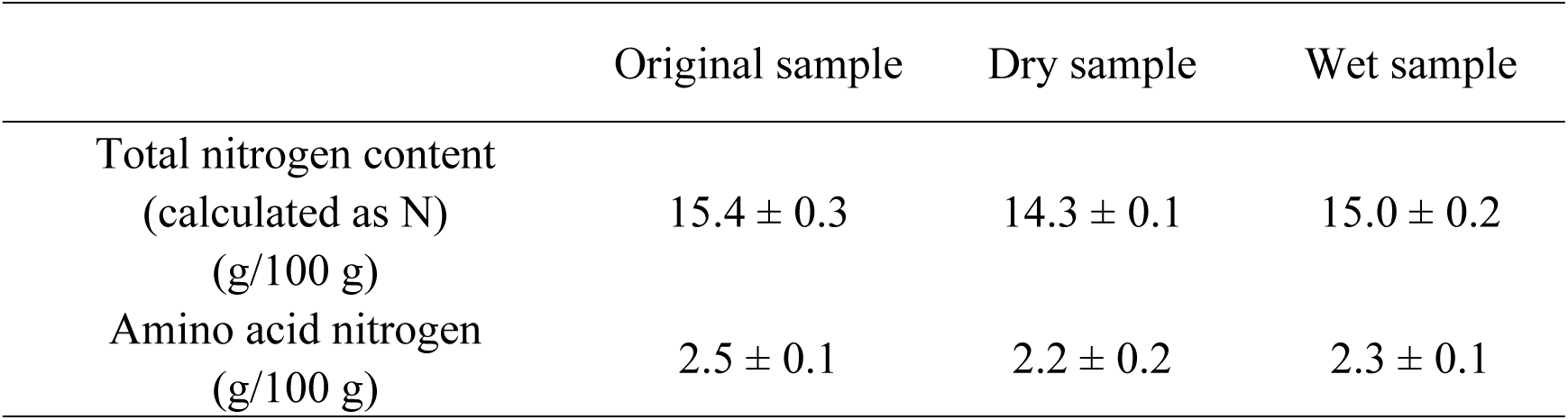
Effects of dry and wet ozonation on total nitrogen and amino nitrogen content of tuna peptide powder.

|  | Original sample | Dry sample | Wet sample |
| --- | --- | --- | --- |
| Total nitrogen content<br>(calculated as N)<br>(g/100 g) | 15.4 ± 0.3 | 14.3 ± 0.1 | 15.0 ± 0.2 |
| Amino acid nitrogen<br>(g/100 g) | 2.5 ± 0.1 | 2.2 ± 0.2 | 2.3 ± 0.1 |

### Low-/High-Temperature Stability Tests of Cosmetics Containing Ozone-treated Tuna Peptides

The qualitative sensory scores (fishy odor/taste) of cosmetic samples containing 1% ozone-treated tuna peptides remained unchanged in the low- and high-temperature stability tests (Table 6). In other words, the fishy odor/taste of the samples did not increase due to changes in temperature. Moreover, the ozone-treated samples were more sensorily acceptable than the untreated sample. This demonstrates the stability of the deodorizing effect of ozonation on tuna peptide powder and supports their use as an ingredient in cosmetics^[26]^. As a result, there is a growing demand for alternative energy sources, and ozone-treated marine animals have become a sustainable and responsible choice.

**Table 6.**
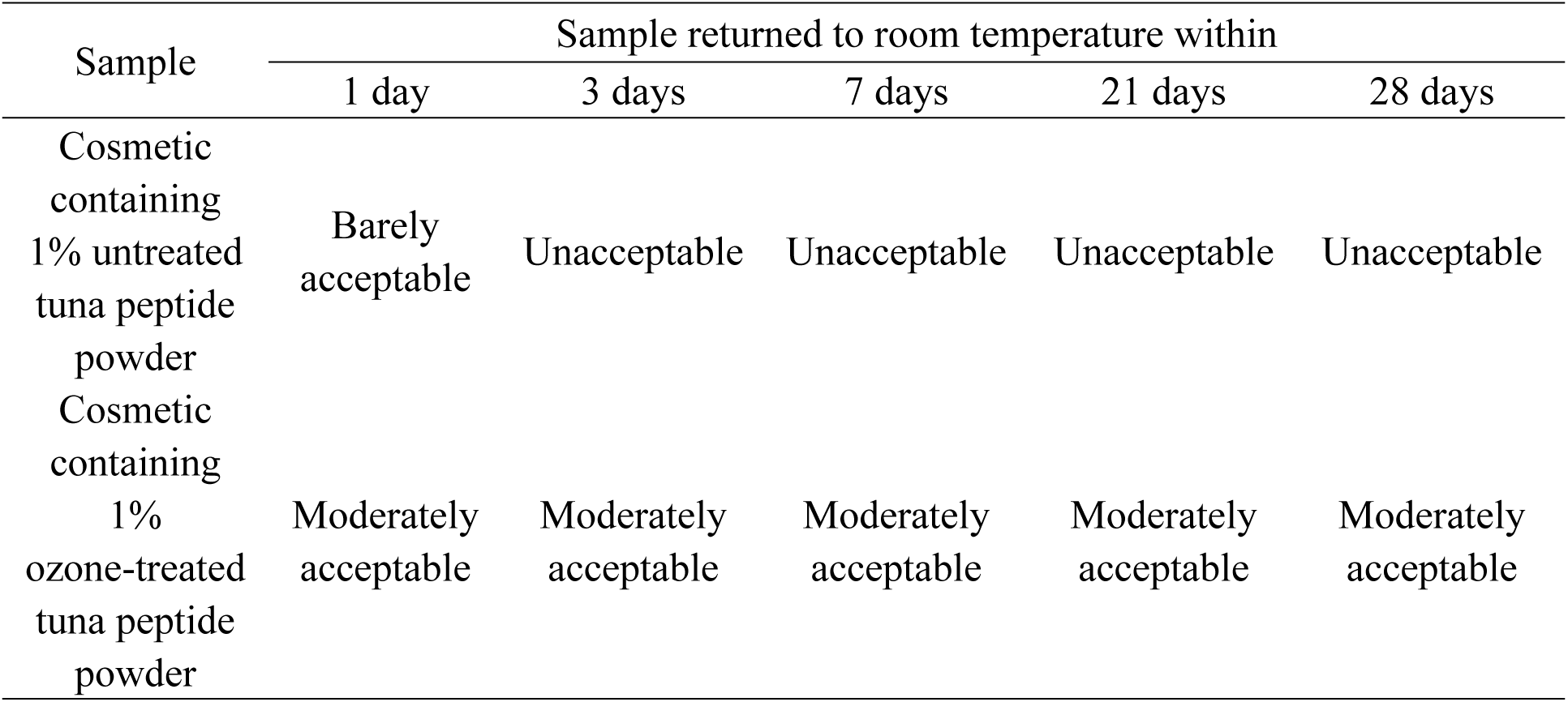
Sensory scores for cosmetic samples during low-/high-temperature stability tests.

## GC-MS RESULTS AND ANALYSIS

The samples treated by wet ozone oxidation with the highest sensory scores were further analyzed by GC-MS. The data in Table 7 were derived from qualitative and quantitative measurements. Among volatile substances with fishy odor, aldehydes have a very low odor threshold. The odor threshold of the main fishy volatile substance, caproaldehyde, is about 0.15–4.3 ppm^[17][27]^. The results showed that the hexaldehyde content decreased by 66.4% after ozone oxidation as they were converted into non-volatile or non-irritating substances. The content of two volatile aldehydes, octyl and non-aldehydes, increased significantly. Manouchehr et al.^[28]^ found that the threshold of volatile aldehydes was very low and that complete addition would occur at low concentrations. Short aldehydes have a pungent odor, while longer aldehydes above C8 have a fruity aroma. The odor threshold of the significantly increased content of these aldehydes is higher, having a non-irritating odor, a lower contribution to the fishy taste, and can even mask the fishy taste. This conclusion is consistent with the reduced fishiness observed in sensory evaluation (Table VII). The mitigation of the fishy taste is essential for its application in the cosmetics industry and offers valuable insights into the high value utilization of marine products.

**Table 7.** Changes in the quality of fishy volatiles before and after oxidation.

| Volatile substance | Original peak area | Peak area of treated sample |
| --- | --- | --- |
| n-Hexaldehyde | 1,566,980 | 525,369 |
| Nonaldehyde | 718,540 | 1,762,707 |
| Octyal | 188,090 | 282,644 |

## DISCUSSION

Ozone is a strong oxidizing agent with a redox potential of 2.07 V, second only to fluorine in oxidizing capacity. As such, it exhibits a very strong oxidizing capacity, which can be used to oxidize and degrade a wide range of organics, including aromatic compounds, unsaturated compounds, difficult-to-biodegrade organics, and hazardous organics with high toxicity^[14]^.

There are two types of ozone oxidation: direct oxidation and indirect oxidation. Dry oxidation operates through an direct mechanism, which typically occurs in solutions containing numerous free radical reaction chain terminators, such as carbonic acid. During direct oxidation, ozone selectively reacts with the organic matter in the peptide powder, resulting in minimal changes in the total organic carbon content before and after oxidation. Ozone can transform part of the organic matter into simpler forms, such as carboxylic acids, or directly oxidize it into carbon dioxide and water.^[15]^ Under direct oxidation conditions, ozone reacts more quickly with unsaturated compounds such as double bonds and aromatic compounds with electron-supplying substituents (phenolic hydroxyl groups).

Wet ozone oxidation operates through an indirect mechanism. In the presence of free radical activators and promoters, ozone produces a large number of free hydroxyl radicals within the reaction system. These hydroxyl radicals undergo a chain reaction, producing more reactive radicals. The reaction rate between these reactive radicals and organic matter is closely aligned with the rate of mass transfer.^[13]^ The hydroxyl radical (-OH) produced in the indirect oxidation process serves as the best oxidant for deep oxidation, which rapidly oxidizes the organic matter and reduces the organic carbon content in water^[14]^.

In addition, O_3_ shows different deodorizing effects, with several factors influencing its removal efficacy^[22][26]^. (1) The amount of O_3_ dosage directly impacts its removal effect. At lower ozone concentrations, the increase in ozone concentration accelerates the reaction rate; however, beyond a certain dosage, further increments do not significantly improve the removal effect, which may be due to the increased concentrations of hydroxyl radicals, collisions between reactive radicals, and heightened chances of ozone resynthesis. (2) Within a certain range, longer contact times lead to more complete reactions, and the removal effect is better than a shorter duration. (3) Different reaction conditions, such as wet and dry environments, yield different effects. In the wet ozone method, the production of more -OH radicals occurs, serving as the best oxidant in advanced oxidation and quickly oxidizing fishy substances in tuna. (4) Temperature also influences the removal effect. It influences the reaction rate of the ozone oxidation system, with an increase in temperature leading to an increase in the reaction rate constant. In wet reactions, heightened temperatures make ozone more likely to escape from water, resulting in a decrease in the ozone concentration; this, in turn, reverses the ozone oxidation reaction equilibrium and affects the reaction speed between ozone and organic matter.

## CONCLUSIONS

The deodorization effect of ozone oxidation on tuna polypeptide was studied, and the optimum deodorization reaction conditions were determined. The results showed that the optimal conditions for dry ozonation were as follows: temperature, 35°C; ozone concentration, 99.1 mg/L; reaction time, 180 min (sensory score 3.4). At an ozone concentration of 99.1 mg/L, a reaction time of 40 min, and a reaction temperature of 50°C, the sensory score of wet ozonation was the lowest (2.3 points). Gas chromatography–mass spectrometry analysis showed that the volatile odor substance, hexanal, decreased by 66.4%, while the content of non-irritating odor aldehydes (non-aldehydes and octanal) with higher odor thresholds increased, reducing the overall odor value of the tuna peptide powder. Ozonation has a significant deodorizing effect on tuna peptide powder, maintaining the protein composition of the tuna peptide powder and retaining its nutritional value. In addition, the low-temperature stability tests showed that the sensory score of ozone-treated tuna peptide powder in cosmetics remained stable and did not increase with changes in temperature. In comparison to other deodorizing methods, ozone deodorization demonstrates greater stability and significant advantages. Among them, the processed tuna peptide powder has a prospective application in the field of cosmetics. Therefore, this study not only provides technical support for the high value utilization of marine protein resources, but also helps to solve the environmental pollution problem of aquatic processing by-products, thereby contributing to the advancement of sustainable blue cosmetics.

## ACKNOWLEDGMENTS

This research was funded by the 2024 Guangdong Province Special Funding for Marine Economy Development (GDNRC[2024]49); the Scientific Research Project of Guangdong Provincial Department of Education (2023KTSCX132); the 2021 Guangdong Provincial Universities Characteristic Innovation Project (2021KTSCX114); and the University-level Scientific Research Project of Foshan University of Science and Technology (CGZ0400166). We thank LetPub (www.letpub.com) for its linguistic assistance during the preparation of this manuscript.

## DISCLOSURE STATEMENT

No potential conflict of interest was reported by the authors.

## Notes

### Competing Interest Statement

The authors have declared no competing interest.

